# Developing *Wolbachia*-based disease interventions for an extreme environment

**DOI:** 10.1101/2022.07.26.501527

**Authors:** Perran A. Ross, Samia Elfekih, Sophie Collier, Melissa J. Klein, Su Shyan Lee, Michael Dunn, Sarah Jackson, Yexin Zhang, Jason K. Axford, Xinyue Gu, Majed S. Nassar, Prasad N. Paradkar, Essam A. Tawfik, Francis M. Jiggins, Abdulaziz M. Almalik, Mohamed B. Al-Fageeh, Ary A. Hoffmann

**Affiliations:** Pest and Environmental Adaptation Research Group, Bio21 Institute and the School of Biosciences, University of Melbourne, Parkville, VIC, Australia; CSIRO Health and Biosecurity, Australian Centre for Disease Preparedness (ACDP), Geelong, VIC, Australia; Department of Genetics, University of Cambridge, Cambridge, United Kingdom; National Center of Biotechnology, Life Science and Environment Research Institute, King Abdulaziz City for Science and Technology (KACST), Riyadh, Saudi Arabia

## Abstract

*Aedes aegypti* mosquitoes carrying self-spreading, virus-blocking *Wolbachia* bacteria are being deployed to suppress dengue transmission. However, there are challenges in applying this technology in extreme environments. We introduced two *Wolbachia* strains into *Ae. aegypti* from Saudi Arabia for a release program in the hot coastal city of Jeddah. *Wolbachia* reduced infection and dissemination of dengue virus (DENV2) in Saudi Arabian mosquitoes and showed complete maternal transmission and cytoplasmic incompatibility. *Wolbachia* reduced mosquito heat tolerance and egg viability, with the *Wolbachia* strains showing differential thermal stability. *Wolbachia* effects were similar across mosquito genetic backgrounds but we found evidence of local adaptation, with Saudi Arabian mosquitoes having lower egg viability but higher adult desiccation tolerance than Australian mosquitoes. Genetic background effects will influence *Wolbachia* invasion dynamics, reinforcing the need to use local genotypes for mosquito release programs, particularly in extreme environments like Jeddah. Our comprehensive characterization of *Wolbachia* strains provides a foundation for *Wolbachia*-based disease interventions in harsh climates.

## 1. Introduction

*Aedes aegypti* mosquitoes are important arbovirus vectors that are widespread throughout tropical and subtropical climates^1^. *Aedes aegypti* have adapted to lay eggs in human-made containers, where they can resist desiccation and remain quiescent for long periods before hatching^2^. Urbanization has contributed to the rapid global spread of *Ae. aegypti* through an increased availability of larval habitats and human hosts, which females feed on to produce eggs^3^. *Aedes aegypti* are capable of rapid adaptive changes in novel environments, with examples including the rapid evolution of resistance to satyrization^4^ and insecticides^5^. While the distribution of *Ae. aegypti* is limited by climate, evolutionary changes will enable *Ae. aegypti* to adapt to new ecological niches and in response to climate change^6, 7, 8^. Globally, *Ae. aegypti* populations show strong genetic differentiation^9, 10^ and there is evidence of local adaptation, particularly for insecticide resistance^11^ and vector competence^12, 13^. Studies of populations collected within a single region show genomic evidence of local adaptation to climate^14^, as well as variation in cold tolerance^15^ and egg survival time^16^ linked to climate. However, the extent of variation in environmental stress tolerance across the global distribution of *Ae. aegypti* is currently unknown^7, 8^.

Many arboviral diseases lack an effective vaccine and their control often relies on the management of vector mosquito populations^17^. In the last decade, releases of mosquitoes carrying self-spreading, virus-blocking strains of *Wolbachia* bacteria have become a leading approach for arbovirus control^18, 19, 20^. *Wolbachia* bacteria occur naturally in about half of all insect species^21, 22, 23^ and have been introduced to *Ae. aegypti* artificially through microinjection^24^. *Wolbachia* infections are transmitted from mother to offspring and can spread through insect populations by providing a relative reproductive advantage to females that carry the infection^25^. *Wolbachia* infections in mosquitoes can induce cytoplasmic incompatibility, where uninfected females that mate with *Wolbachia-* infected males do not produce viable offspring^26^. When *Wolbachia*-infected mosquitoes are released into the environment, cytoplasmic incompatibility helps to drive the infection into the mosquito population. Most *Wolbachia* transinfections in *Ae. aegypti* also reduce arbovirus replication and transmission by mosquitoes^27, 28, 29, 30^, making the establishment of *Wolbachia*-infected mosquitoes an effective way to reduce the local spread of disease. *Wolbachia* strains that induce cytoplasmic incompatibility can also be used to suppress mosquito populations through male-only releases^31, 32^.

*Wolbachia* ‘population replacement’ programs are now operating with the *w*Mel *Wolbachia* strain in over 10 countries (https://www.worldmosquitoprogram.org/en/global-progress) and the *w*AlbB strain in Kuala Lumpur, Malaysia^19^. The establishment of *Wolbachia* infections at high frequencies substantially reduces arbovirus transmission^19, 20, 33, 34^. However, the ability of *Wolbachia* infections to stably establish in *Ae. aegypti* populations depends on the choice of *Wolbachia* strain and local environmental conditions. In Rio de Janeiro, Brazil, *w*Mel initially failed to establish because the released mosquitoes lacked local insecticide resistance alleles^35, 36^. *Wolbachia* establishment may also be limited by climate and the types of larval habitats available. For the *w*Mel *Wolbachia* strain which is vulnerable to high temperatures^30, 37, 38^, seasonal fluctuations in *w*Mel infection frequencies occur in some locations^39, 40^, which may reflect high temperatures in larval habitats^41^. In Malaysia, the establishment of *w*AlbB varied substantially between release sites^19^, likely due to high host fitness costs of this strain in quiescent eggs^42, 43^.

When performing a *Wolbachia* release program it is important to understand the effects of *Wolbachia* infections in a local context. Release strains are usually characterized under laboratory or semi-field conditions prior to release^30, 44, 45, 46, 47^, but these conditions may not accurately reflect the situation in the field. *Wolbachia* infections in *Ae. aegypti* induce host fitness costs under sub-optimal conditions, reducing the viability of quiescent eggs^42, 48, 49^, thermal tolerance^41, 50^, starvation tolerance^51, 52^ and performance when feeding on non-human blood^53, 54, 55^. The impact of these host fitness costs on the ability of *Wolbachia* to establish is likely to depend on local environmental conditions; locations that experience long dry seasons, extreme temperatures or intense larval competition could make *Wolbachia* establishment more challenging. Given that local adaptation can also lead to differences in stress tolerance^56^, it is plausible that *Wolbachia* infections could interact with mosquito genotype.

Releases of *Ae. aegypti* with *Wolbachia* strains are underway in Jeddah, Saudi Arabia for dengue control and preparatory work has been initiated to characterize the local mosquito population^57, 58^. The city of Jeddah has a climate with low rainfall (BWh; Hot Desert Climate) under the Koppen’s classification. It features a tropical and subtropical temperature range^59^, with a yearly average high temperature of 35°C and an average low temperature of 23°C (reaching an average high of 39°C in July and an average low of 20°C in January). The yearly average relative humidity ranges between 60 to 67%, and rainfall is 60mm on average per year^60^. Rainfalls in Jeddah are scattered, and the rainy season usually occur between November and January^61^. Insecticides are used widely for mosquito management in the region which has led to a high prevalence of insecticide resistance^57, 62^.

In this study, we developed two *Ae. aegypti* strains for the Saudi Arabia *Wolbachia* release program. We introduced the *w*AlbB^63^ and *w*MelM^64^ infections to locally-sourced mosquitoes through backcrossing then compared their performance to counterparts from Cairns, Australia. We show that both strains provide effective virus blocking in Saudi Arabian mosquitoes. We then performed a comprehensive phenotypic characterization of these mosquito populations with a focus on desiccation, storage and heat stress tolerance; these traits are likely to be important given that Jeddah experiences low rainfall and hot summer temperatures. We identified novel effects of *Wolbachia* infections on thermal and desiccation tolerance and found clear differences between Australian and Saudi Arabian populations indicative of local adaptation. Our results will help to inform *Wolbachia* release programs in this environment and similar climatic scenarios.

## 2. Results

### 2.1 Generation of stable Wolbachia strains in Saudi Arabian mosquitoes

We generated mosquito populations with two genetic backgrounds (Saudi Arabia, S and Australia, Au) and three *Wolbachia* infection types (*w*AlbB, *w*MelM and Uninfected) through microinjection^63, 64^, followed by repeated backcrossing of *Wolbachia*-infected females to wild-caught males from Cairns, Australia or Jeddah, Saudi Arabia. Both the *w*AlbB and *w*MelM infections showed complete maternal transmission in Saudi Arabian mosquitoes, with 100/100 progeny from ten *Wolbachia-* infected mothers for each strain testing positive for *Wolbachia* by PCR (lower 95% confidence interval: 0.963). This was corroborated in a second experiment with *w*AlbB S showing complete maternal transmission to 75 progeny from 17 *Wolbachia*-infected mothers (lower 95% confidence interval: 0.952). These results are consistent with previous work on these strains in an Australian background^63, 64^.

We then tested the ability of *w*AlbB to induce cytoplasmic incompatibility in Saudi Arabian mosquitoes. *w*AlbB-infected males induced complete cytoplasmic incompatibility with uninfected females, with no eggs hatching (Table 1). All other crosses were fertile, but egg hatch proportions were lower in crosses with *w*AlbB-infected females, consistent with later experiments (see below).

**Table 1.** Cytoplasmic incompatibility induction by *w*AlbB-infected males in *Aedes aegypti* from a Saudi Arabian background. Median egg hatch proportions and confidence intervals (lower 95%, upper 95%) are shown.

| Male population | Female population |  |
| --- | --- | --- |
|  | Uninfected S | wAlbB S |
| Uninfected S | 0.965 (0.920, 0.986) | 0.833 (0.661, 0.926) |
| wAlbB S | 0 (0, 0) | 0.893 (0.800, 0.963) |

To test for potential changes in the *Wolbachia* genome in Saudi Arabian mosquitoes, the *Wolbachia* genomes of the *w*AlbB Au and *w*AlbB S populations were Illumina-sequenced, resulting in coverage of 91X for the genome in the Saudi Arabian genetic background and 143X in the Australian genetic background. The two genomes are identical, both differing from the reference genome by four single nucleotide variants and one insertion (Table S1). These results point to the stability of the *w*AlbB genome across time and in different mosquito genetic backgrounds.

### 2.2 *Wolbachia infections block DENV2 replication and dissemination in* Ae. aegypti *with a Saudi Arabian background*

We previously demonstrated that *w*AlbB and *w*MelM reduce DENV2 virus titers in Australian *Ae. aegypti^64^.* Here, we estimated the effects of these *Wolbachia* strains on DENV2 blocking in *Ae. aegypti* from a Saudi Arabian background. Both *w*AlbB S and *w*MelM S showed reduced midgut (Two-tailed Fisher’s exact test, *w*AlbB: P = 0.025, *w*MelM: P < 0.001, Figure 1A) and carcass (*w*AlbB: P < 0.001, *w*MelM: P < 0.001, Figure 1B) DENV2 infection frequencies compared to Uninfected S. For samples that tested positive, *w*AlbB S had fewer DENV2 copies compared to Uninfected S in midgut samples (Mann-Whitney U: Z = 4.431, P < 0.001). DENV2 blocking appeared to be particularly strong for *w*MelM, with reduced midgut infection frequencies compared to *w*AlbB (Two-tailed Fisher’s exact test, P < 0.001). Near-complete DENV2 blocking by *w*MelM was further demonstrated in an additional comparison of the *w*MelM S and Uninfected S populations (Figure S1). We were unable to perform other comparisons of DENV2 copy numbers owing to the very low infection frequencies in *Wolbachia*-infected mosquitoes as noted above. We also had low power to estimate effects of *Wolbachia* infection on DENV2 transmission due to low saliva infection frequencies in the uninfected population (13.3% positive, Figure 1C), however all *w*AlbB S and *w*MelM S saliva samples were negative for DENV2. In a separate experiment, *w*AlbB also showed blocking in an Australian background, with *w*AlbB Au having fewer DENV2 copies in the midgut (Z = 2.748, P = 0.006) and carcass (Z = 4.935, P < 0.001) compared to Uninfected Au (Figure S2). Together, these results demonstrate DENV2 blocking of both *w*AlbB and *w*MelM in Saudi Arabian *Ae. aegypti.* Saliva infection assays likely underrepresent transmission potential^65^, but our analyses of midgut and carcass infections suggest that *Wolbachia* infections will likely reduce DENV2 transmission.

**Figure 1.**
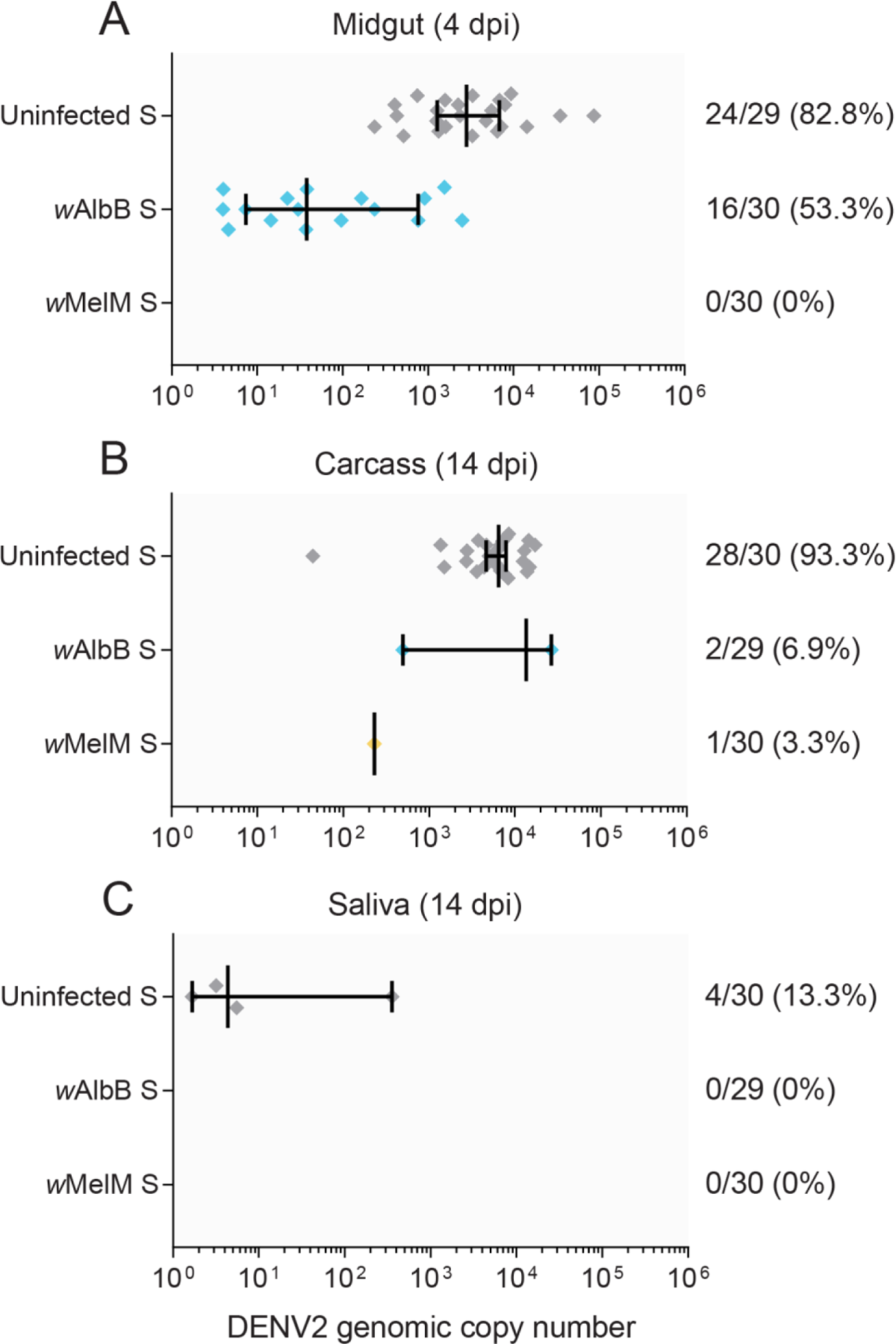
Comparison of DENV2 infection frequencies and copy numbers in uninfected, *w*AlbB-infected and *w*MelM-infected *Ae. aegypti* with a Saudi Arabian background. We measured (A) midgut infection at 4 dpi, (B) dissemination to the carcass at 14 dpi and (C) saliva infection at 14 dpi as a proxy for transmission. Each symbol shows data from a single DENV2-positive sample (negative samples were excluded), while vertical lines and error bars represent medians and 95% confidence intervals. Numbers to the right of indicate the number of individuals positive for DENV2 out of the total number tested, followed by the percent positive.

### 2.3 *Rapid proliferation of wAlbB* Wolbachia *in adult mosquitoes*

*Wolbachia* genotypes that reach a high density in their insect hosts tend to block viral infection more strongly but impose greater fitness costs on their hosts. We found a significant effect of adult age on *w*AlbB density (GLM: F_5,236_ = 25.163, P < 0.001), with median density increasing ~17-fold in the 35 days after adult females eclosed (Figure S3A). The change in density was greatest in the first week post-eclosion, where there was a ~6-fold increase in *Wolbachia* density.

Heterogeneity within vector populations can impact disease transmission^66^. Within a single time point there is a considerable variation between individuals in *Wolbachia* density—on average the mosquito with the highest density had 632 times higher *Wolbachia* density than the individual with the lowest density. Furthermore, little of this variation is caused by measurement error. We measured the *Wolbachia* density of each sample twice independently, and within each day 94% percent of the variance in *Wolbachia* density was caused by biological differences between mosquitoes rather than technical variation (repeatability=0.94; Figure S3B).

### 2.4 Mosquito background influences body size but not larval development

We measured larval development time, survival to pupa, sex ratio and adult wing length for each *Wolbachia* infection type across the Australian and Saudi Arabian mosquito backgrounds. We found no significant effect of *Wolbachia* infection type for any trait tested when significance thresholds were adjusted for Bonferroni correction(GLM: all P > 0.012, adjusted α: 0.008 given that six traits were scored). However, wing length was strongly influenced by background in both females (F_1,130_ = 41.378, P < 0.001) and males (F_1,122_ = 76.234, P < 0.001), with *Ae. aegypti* with a Saudi Arabian background being consistently larger than *Ae. aegypti* with an Australian background (Table 2).

**Table 2.** Larval development and size of *Wolbachia*-infected and uninfected *Aedes aegypti* from Australian and Saudi Arabian backgrounds. Medians and confidence intervals (lower 95%, upper 95%) are shown for each trait.

| Population | Proportion survival to pupa | Larval development time (days) |  | Sex ratio (males/total) | Wing length (mm) |  |
| --- | --- | --- | --- | --- | --- | --- |
|  |  | Males | Females |  | Males | Females |
| Uninfected Au | 0.99 (0.97, 1.00) | 6.38 (6.33, 6.83) | 7.05 (6.83, 7.15) | 0.47 (0.39, 0.60) | 2.12 (2.07, 2.19) | 2.79 (2.73, 2.83) |
| Uninfected S | 0.93 (0.91, 0.99) | 6.40 (6.14, 7.01) | 6.98 (6.90, 7.28) | 0.55 (0.42, 0.62) | 2.22 (2.18, 2.24) | 2.90 (2.87, 2.92) |
| wAlbB Au | 0.98 (0.95, 1.00) | 6.62 (6.50, 6.74) | 7.04 (6.79, 7.19) | 0.48 (0.43, 0.51) | 2.14 (2.13, 2.16) | 2.75 (2.65, 2.85) |
| wAlbB S | 0.97 (0.94, 0.99) | 6.54 (6.33, 7.08) | 7.09 (6.81, 7.50) | 0.50 (0.45, 0.52) | 2.24 (2.22, 2.29) | 2.85 (2.72, 2.95) |
| wMelM Au | 0.97 (0.90, 1.00) | 6.66 (6.34, 6.88) | 7.10 (6.80, 7.17) | 0.49 (0.46, 0.52) | 2.12 (2.08, 2.17) | 2.72 (2.70, 2.80) |
| wMelM S | 1.00 (0.95, 1.00) | 6.65 (6.50, 6.81) | 7.19 (7.05, 7.50) | 0.51 (0.47, 0.58) | 2.23 (2.20, 2.29) | 2.86 (2.84, 2.92) |

### 2.5 Severe cumulative fitness costs of wAlbB infection following egg storage

We measured the viability of *w*AlbB-infected and uninfected eggs from Australian and Saudi Arabian backgrounds following storage. Egg viability decreased across time, particularly in Saudi Arabian populations in the early time points, where egg hatch proportions were below 0.5 at week 5, but costs of *w*AlbB infection also became apparent in later time points (Figure 2A). Both *Wolbachia* infection (GLM: all P < 0.014) and background (all P < 0.001) had significant effects on (logit transformed) egg hatch proportions at each time point. At week 11, we also found an interaction between *Wolbachia* infection and background (F_1,44_ = 23.428, P < 0.001), where the effects of *w*AlbB infection were more severe in the Australian background.

**Figure 2.**
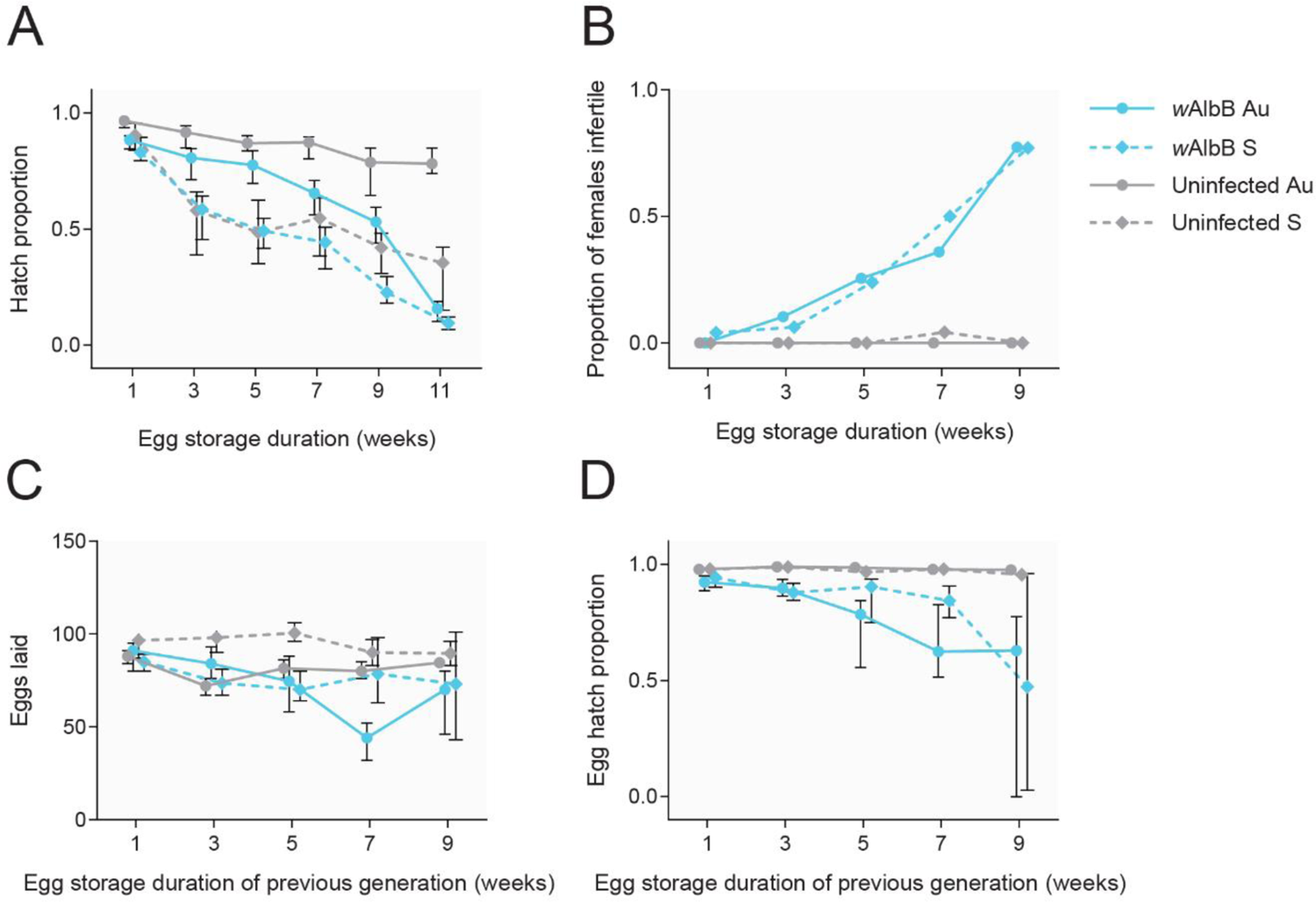
Cross-generational effects of egg storage on *w*AlbB-infected and uninfected *Ae. aegypti* from Australian and Saudi Arabian backgrounds. (A) Egg hatch proportions of eggs stored for 1 to 11 weeks. Females hatching from eggs stored for different periods were then reared to adulthood and measured for their fertility in panels B-D. (B) Proportion of females hatching from stored eggs that were infertile (with no eggs laid) following a blood meal. (C) Number of eggs laid by fertile females hatching from stored eggs in the previous generation. (D) Egg hatch proportions of fertile females hatching from stored eggs in the previous generation. Symbols show medians, while error bars are 95% confidence intervals. Error bars in panel B are not shown as data are binomial. Data points within each temperature cycle have been offset to avoid overlap.

We then measured the fertility of females hatching from stored eggs. *w*AlbB-infected females stored as eggs became increasing infertile with storage time (Figure 2B). After 9 weeks of storage, 77%

(binomial confidence interval: 63-87%) of females from the *w*AlbB Au and *w*AlbB S populations were infertile (with no eggs laid). In contrast, females from uninfected populations remained fertile, with no infertile females at week 9 (binomial confidence interval: 0-7%). The fecundity and (logit transformed) egg hatch proportion of fertile females stored as eggs for 1 week was influenced by *w*AlbB infection (GLM: fecundity: F_1,190_ = 7.369, P = 0.007, egg hatch: F_1,190_ = 39.279, P < 0.001) but not background (fecundity: F_1,190_ = 0.238, P = 0.626, egg hatch: F_1,190_ = 0.319, P = 0.573). We found similar effects after 9 weeks of storage, but with a stronger effect of *w*AlbB infection on egg hatch (F_1,117_ = 124.292, P < 0.001). The egg hatch of uninfected populations remained above 95% across all time points, but this declined to 50-60% in both *w*AlbB-infected populations (Figure 2D). In a scenario where eggs are stored for 9 weeks, *w*AlbB-infected populations had a relative fitness of 8.15% (Australian background) and 4.99% (Saudi Arabian background) compared to their uninfected counterparts when considering effects across two generations.

### 2.6 Genetic background influences adult desiccation tolerance and water loss

We measured adult mosquito survival at low (~15%) and high (~80%) RH as an estimate of desiccation tolerance. Females survived longer than males and we therefore analyzed data separately for each sex. In the low RH experiment (Figure 3A-B), we observed significant effects of background (Cox regression: females: χ^2^ = 28.088, d.f. = 1, P < 0.001, males: χ^2^ = 22.961, d.f. = 1, P < 0.001) but not *w*AlbB infection (females: χ^2^ = 2.855, d.f. = 1, P = 0.091, males: χ^2^ = 3.444, d.f. = 1, P = 0.063). Similarly, in the high RH experiment (Figure 3C-D), we found effects significant effects of background (Cox regression: females: χ^2^ = 9.397, d.f. = 1, P = 0.002, males: χ^2^ = 9.544, d.f. = 1, P = 0.002) but not *w*AlbB infection (females: χ^2^ = 0.001, d.f. = 1, P = 0.977, males: χ^2^ = 0.304, d.f. = 1, P = 0.582). In both cases, mosquitoes from a Saudi Arabian background had longer survival times compared to mosquitoes from an Australian background, with no clear effect of *w*AlbB infection (Figure 3).

**Figure 3.**
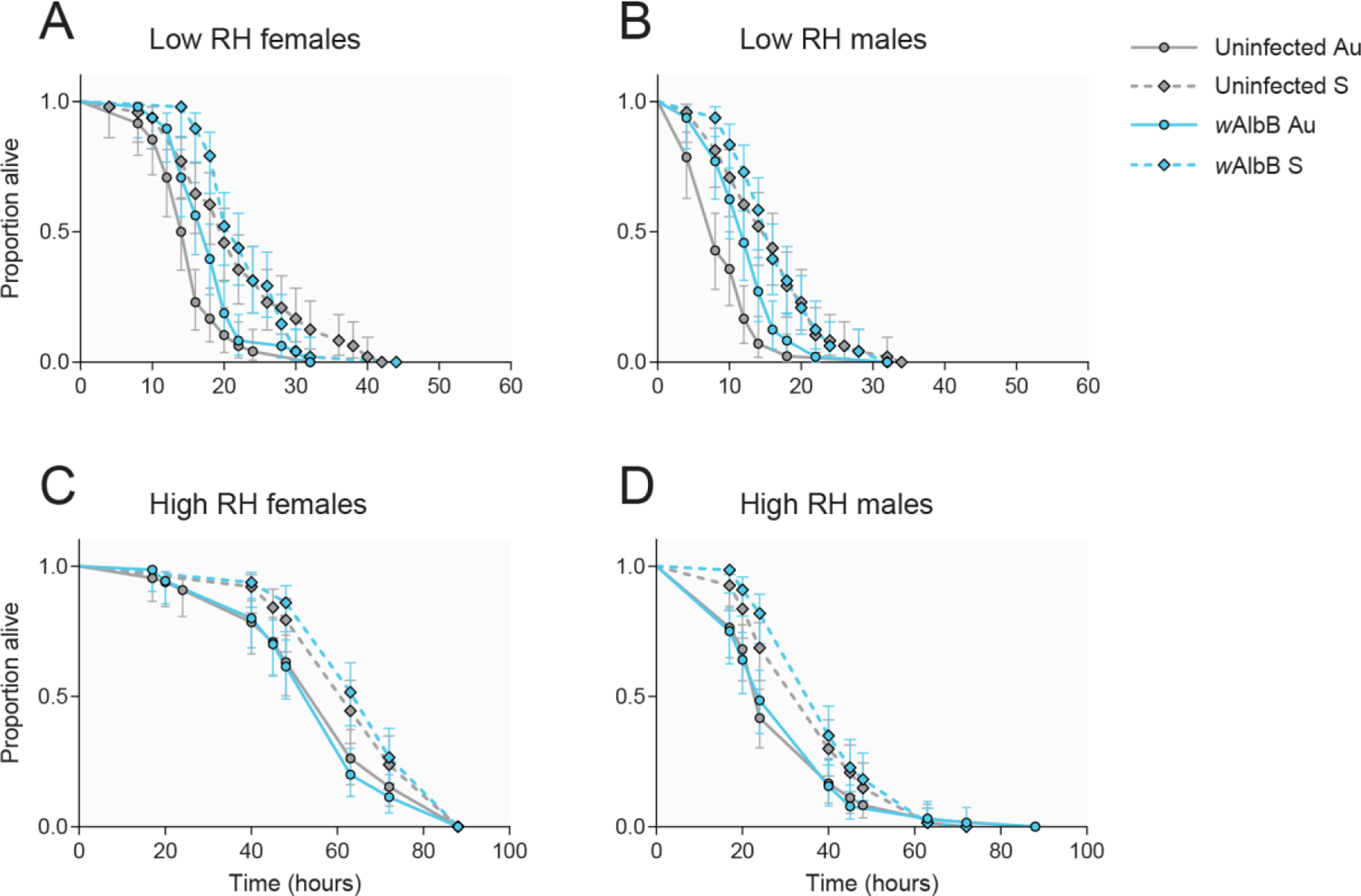
Survival of *w*AlbB-infected and uninfected *Aedes aegypti* adults from Australian and Saudi Arabian backgrounds under low (A-B) and high (C-D) relative humidity. Data are presented separately for females (A, C) and males (B, D). Lines show data from 48 (A-B) or 64 (C-D) adults per population. Error bars represent 95% confidence intervals.

In a separate experiment, we estimated water loss by weighing mosquitoes before and after an 8 hr period at low RH. We found significant effects of background on water loss (expressed as a percentage of water mass lost per hr) in males (GLM: F_1,27_ = 25.357, P < 0.001) but not females (F_1,27_ = 3.035, P = 0.093). Males with an Australian background had a ~50% higher rate of water loss compared to males from the Saudi Arabian background (Table 3). The same pattern was observed for females, but differences were marginal, at ~6% higher.

**Table 3.** Average mass and water loss in *w*AlbB-infected and uninfected *Aedes aegypti* adults from Australian and Saudi Arabian backgrounds under low relative humidity. Medians and standard deviations are shown for each trait.

| Population | Sex | Replicate vials (n = 8 per vial) | Fresh mass per mosquito (mg) | Dry mass per mosquito (mg) | Water mass per mosquito (mg) | Water content (%) | Water loss per hr (% of total water mass) |
| --- | --- | --- | --- | --- | --- | --- | --- |
| Uninfected Au | Female | 8 | 2.10 ± 0.25 | 0.46 ± 0.08 | 1.69 ± 0.27 | 77.43 ± 4.53 | 2.46 ± 1.13 |
| Uninfected S | Female | 7 | 2.07 ± 0.12 | 0.46 ± 0.05 | 1.58 ± 0.09 | 76.03 ± 1.84 | 2.30 ± 0.39 |
| *wAlbB* Au | Female | 8 | 2.64 ± 0.21 | 0.54 ± 0.06 | 2.07 ± 0.17 | 79.10 ± 1.72 | 2.41 ± 0.39 |
| *wAlbB* S | Female | 8 | 1.87 ± 0.30 | 0.37 ± 0.10 | 1.52 ± 0.23 | 81.52 ± 3.16 | 2.28 ± 0.30 |
| Uninfected Au | Male | 7 | 0.93 ± 0.15 | 0.13 ± 0.02 | 0.80 ± 0.15 | 85.74 ± 2.40 | 3.60 ± 1.08 |
| Uninfected S | Male | 8 | 1.00 ± 0.05 | 0.15 ± 0.03 | 0.86 ± 0.05 | 84.71 ± 2.84 | 2.19 ± 0.40 |
| *wAlbB* Au | Male | 8 | 1.17 ± 0.09 | 0.11 ± 0.07 | 1.07 ± 0.09 | 90.51 ± 5.38 | 3.36 ± 0.44 |
| *wAlbB* S | Male | 8 | 1.07 ± 0.27 | 0.16 ± 0.08 | 0.93 ± 0.20 | 85.31 ± 4.39 | 2.25 ± 0.50 |

### 2.7 Wolbachia infection reduces adult heat tolerance and egg survival

We measured the effect of *w*AlbB infection and background on heat tolerance in adult females using a static knockdown assay at 42°C. Knockdown time was highly variable and the relative performance of populations appeared to differ between trials, with a significant interaction between trial and population (GLM: F_15,216_ = 2.949, P < 0.001). Overall, *w*AlbB Au females had a ~30% faster median knockdown time compared to the other three populations (Figure S4). When trials were considered separately, we found significant effects of *w*AlbB infection (Trials 1-6: P = 0.002, 0.983, 0.165, 0.424, 0.964, 0.005) and background (Trials 1-6: P = 0.190, 0.453, 0.038, 0.925, 0.818, 0.001) in only 2/6 trials each, indicating a lack of consistent pattern between trials (Figure 4).

**Figure 4.**
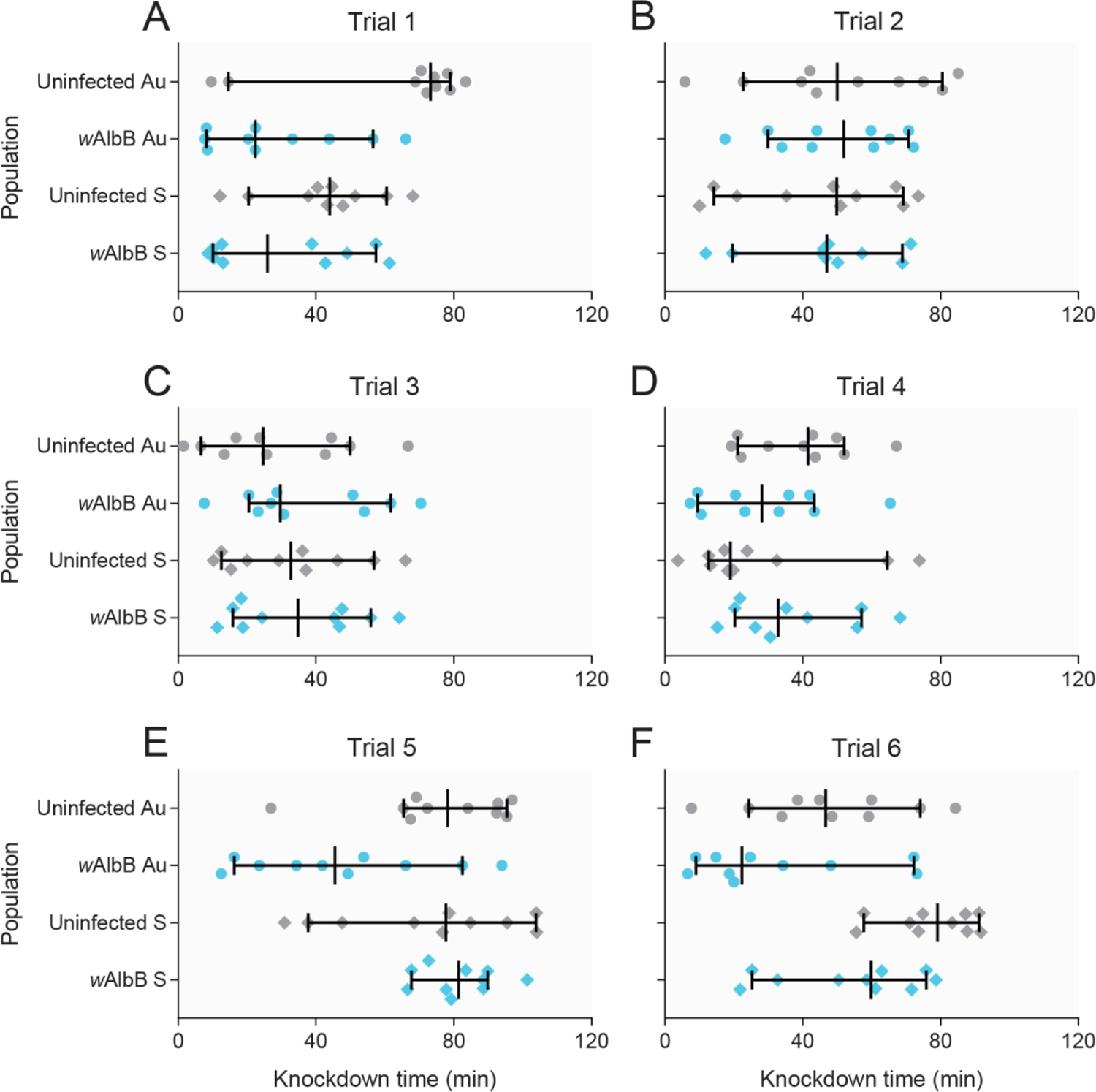
Knockdown time of *w*AlbB-infected and uninfected *Aedes aegypti* females from Australian and Saudi Arabian backgrounds exposed to a 42°C heat shock. Data are shown separately for six individual trials (A-F). Each symbol represents data from a single female, while vertical lines and error bars represent medians and 95% confidence intervals.

We then tested heat tolerance in eggs by exposing them to a range of cyclical temperatures for one week. Egg hatch proportions declined in all populations as temperatures increased, with near-zero hatching when eggs were exposed to a 32-42°C for one week (Figure 5A-B). Egg hatch proportions were lower in *Wolbachia*-infected populations compared to uninfected populations at most high temperatures, suggesting an effect of *Wolbachia* on egg thermal tolerance (Figure 5A-B). However, we also found significant effects of *Wolbachia* infection (GLM: F_2,99_ = 10.305, P < 0.001) and background (F_1,99_ = 9.411, P = 0.003) on (logit) egg hatch proportion in the 26°C control temperature, indicating that *Wolbachia* infection had negative effects independent of temperature. We also found an effect of background in the 31-41°C treatment (F_1,97_ = 11.074, P = 0.001), suggesting that eggs from the Saudi Arabian background had increased tolerance to temperatures near their upper thermal limit.

**Figure 5.**
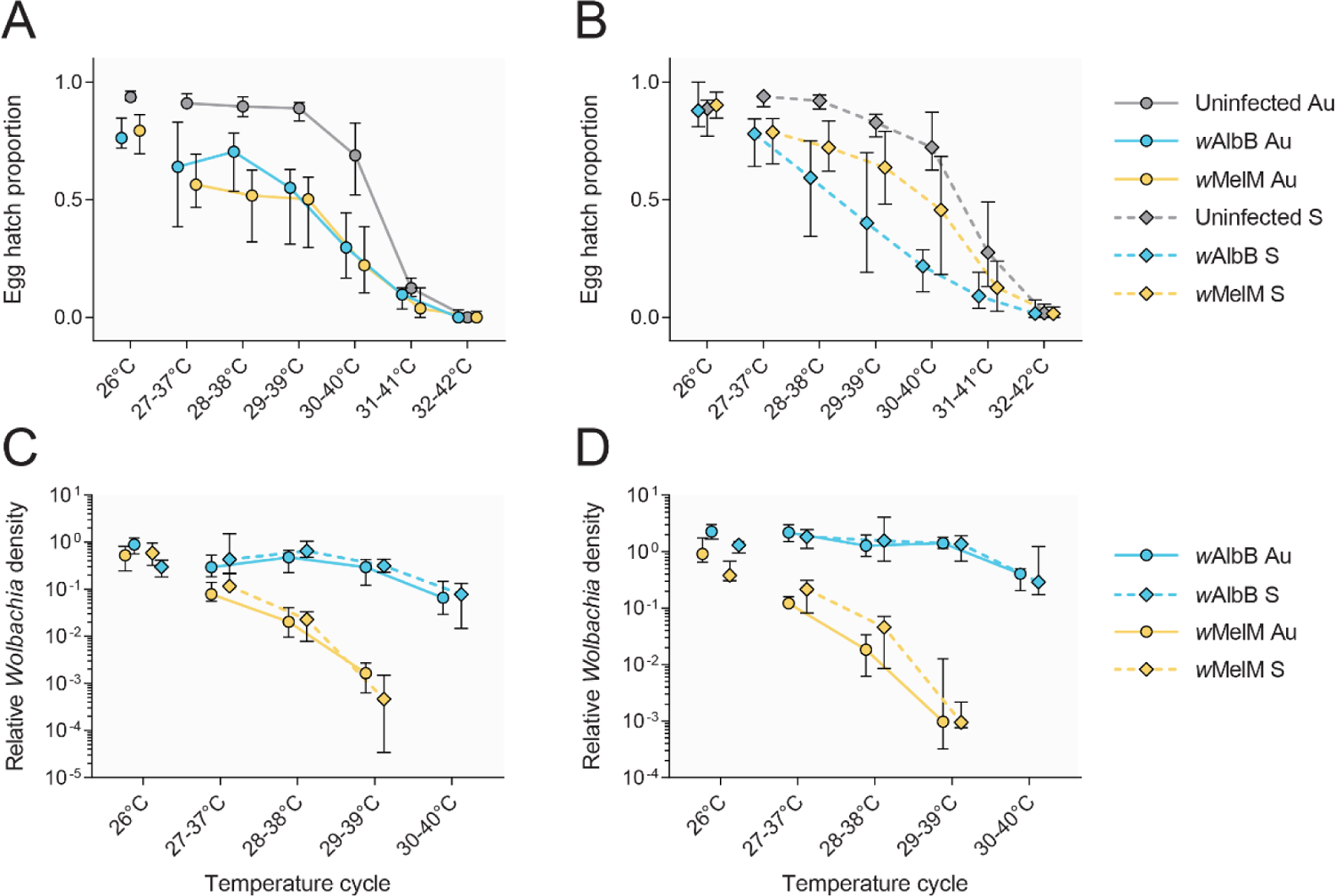
Effects of high cyclical temperatures on egg hatch and *Wolbachia* density in *Wolbachia-* infected and uninfected *Ae. aegypti* from Australian and Saudi Arabian backgrounds. Eggs were placed in thermocyclers set to 26°C (control) or daily cycling temperatures for one week before hatching. Egg hatch proportions are presented separately for Australian (A) and Saudi Arabian (B) populations, with *Wolbachia* density presented separately for (C) females and (D) males. Symbols show medians, while error bars are 95% confidence intervals. Data points within each temperature cycle have been offset to avoid overlap.

We then measured *Wolbachia* density in adults emerging from eggs exposed to heat stress (Figure 5C-D). We found strong overall effects of *Wolbachia* strain (GLM: females: F_1,137_ = 417.422, P < 0.001, males: F_1,138_ = 627.602, P < 0.001) and temperature (females: F_4,137_ = 111.474, P < 0.001 males: F_4,138_ = 76.812, P < 0.001) on *Wolbachia* density, with an interaction between *Wolbachia* strain and temperature (females: F_3,137_ = 92.940, P < 0.001, males: F_3,138_ = 71.569, P < 0.001). *Wolbachia* densities of both strains decreased with increasing temperature, but the decline was much sharper for *w*MelM (Figure 5C-D). We observed complete loss of *w*MelM in some individuals at high temperatures, with 13/36 individuals exposed to 29-39°C testing negative and 34/36 negative at 30-40°C across males and females from both backgrounds. In contrast, all individuals from *w*AlbB-infected populations were positive for *Wolbachia* across all treatments. We found no overall effect of background (females: F_1,137_ = 0.460, P = 0.499, males: F_1,138_ = 0.014, P = 0.907), with *w*MelM and *w*AlbB densities decreasing to a similar extent with temperature both the Australian and Saudi Arabian backgrounds (Figure 5C-D).

We then tested whether the effects of temperature and wAlbB infection on egg hatch were modulated by RH. Eggs from the *w*AlbB S and Uninfected S populations were stored for 3 weeks under two temperature (20-30°C and 29-39°C) and two RH (80% and 20%) conditions. We found significant effects of both temperature (GLM: F_1,139_ = 88.702, P < 0.001) and RH (F_1,139_ = 39.703, P < 0.001) on egg hatch. There was also an interaction between temperature and RH (F_1,139_ = 26.117, P < 0.001), where no eggs hatched under high temperature, low humidity conditions (Figure S5B). Egg hatch proportions were lower in the *w*AlbB S population compared to Uninfected S in all treatments where eggs survived (Figure S5B), however there was no significant effect of *w*AlbB infection overall (F_1,139_ = 2.054, P = 0.154). The reduction in egg hatch caused by *w*AlbB infection was not significantly affected by temperature (population by temperature interaction: F_1,139_ = 0.113, P = 0.738 or humidity (population by humidity interaction: F_1,139_ = 0.028, P = 0.868). *w*AlbB infection also reduced egg hatch in the controls where eggs were not stored (F_1,73_ = 11.779, P < 0.001, Figure S5), demonstrating costs of infection regardless of storage and environmental conditions.

## 3. Discussion

With *Wolbachia* releases taking place in different climate regions, it is important to understand variation in mosquito tolerance to environmental stress and any interactions with *Wolbachia* infections. In this study, we developed and characterized two *Wolbachia* strains for use in population replacement programs under the extreme conditions of Jeddah, Saudi Arabia. Both strains are likely to be suitable for field release due to their strong virus blocking, maternal transmission, incompatibility effects and other characteristics. The *w*MelM strain in particular exhibited very strong blocking in the Saudi Arabian background, making this strain a good candidate for release given that it does not have severe fitness costs^64^. Note that although we did not test *w*MelM rigorously for cytoplasmic incompatibility here, it causes complete cytoplasmic incompatibility in an Australian background^64^ and preliminary work suggests complete cytoplasmic incompatibility in a Saudi Arabian background (with no eggs hatching from >3000 eggs produced from a cross between a group of *w*MelM S males and Uninfected S females).

Our results highlight phenotypic differences between *Ae. aegypti* with genetic backgrounds from Cairns, Australia and Jeddah, Saudi Arabia. The Saudi Arabian mosquitoes showed a higher fecundity and increased adult desiccation tolerance compared to Australian mosquitoes, but the ability of their eggs to withstand long periods of quiescence was surprisingly relatively poor. This geographic variation in stress tolerance may reflect local adaptation to climate^67, 68^ and the type of larval habitats that are common in these locations. Perhaps the unexpected lower quiescent egg viability in Saudi Arabian mosquitoes suggests that eggs are not exposed to long dry periods, despite the very low annual rainfall in this city. The *Ae. aegypti* in this location may use permanent sources of water for oviposition such as underground car parks where we have found evidence of mosquito breeding even in the middle of the dry and hot summer period (S. Elfekih, unpublished observations). In these locations, temperatures can be relatively cool (<30°C) even when ambient temperatures at the surface exceed 40°C. There is very little shade in Jeddah residential areas outside of buildings due to low vegetation cover and this would make it unlikely that quiescent eggs would survive in summer^69^. In contrast, in Cairns there is shade around buildings provided by lush vegetation and across the dry winter period survival of quiescent eggs can be high^69^.

The increased desiccation tolerance in Saudi Arabian mosquitoes suggests that adult mosquitoes are adapted to low RH conditions. In Cairns, relative humidity averages 63% and the humidity peaks in the summer wet season (www.bom.gov.au) when mosquito numbers are highest but average temperatures are still below 30°C. In Jeddah, the average humidity is similar but is around 60% when *Ae. aegypti* numbers are high in February-March which coincides with higher average temperatures of 30°C or more. Adult insect desiccation stress will likely depend on a combination of temperature and humidity at times when the adults are active^70^ and combined temperature/humidity metrics such as the drying power of air are often used in assessing insect species differences in desiccation resistance^71^. We would therefore expect the adults from Saudi Arabia to be exposed to additional desiccation stress as they move about beyond built structures; based on genomic data we have previously found that *Ae. aegypti* do move tens or even hundreds of meters across a generation although this can be limited by the presence of wide roads^58^.

The phenotypic variation that exists in *Ae. aegypti* across its current distribution is still poorly understood given that large scale comparisons across geographic regions have not been completed, but there are indications that population variation exists for several traits^11, 12, 15, 16^. In other Diptera like various *Drosophila* species, geographic variation in traits has been linked to climatic factors^72, 73^. There is also a rich literature on climate adaptation in insect populations associated with quiescence and diapause at various developmental stages^74, 75^ which have developed in many mosquito species where these phenotypes aid in population persistence and spread^2^. Our data suggest that it may be worthwhile assessing the relative desiccation resistance of different developmental stages of *Ae. aegypti* and linking these not only to climate but also to the nature of breeding sites present in different locations that affect the dynamics of the local populations.

Endosymbionts can modulate the desiccation^76, 77^ and thermal^78, 79^ tolerance of their hosts. Hence, we investigated the phenotypic effects of *Wolbachia* under a range of stressful environmental conditions. We found clear costs of *Wolbachia* infection to several traits, including egg hatch, particularly in quiescent eggs, reduced adult thermal tolerance for *w*AlbB, and greatly reduced female fertility. These costs were largely independent of genetic background, but their importance will depend on background and environment. As demonstrated previously^43^, effects of *Wolbachia* on viability and fertility after egg storage are substantial, but may be less relevant in areas with continuously available larval habitats or in locally-adapted mosquitoes where egg viability is already poor. Given that *Wolbachia* establishment can vary substantially between locations^34, 40^, even in similar climates, the local environment and mosquito adaptation to this environment are likely to be key drivers of *Wolbachia* establishment success.

We investigated novel traits, adult desiccation tolerance and water loss, in this study and found no clear effect of *Wolbachia* on either trait. Consistent with a recent study, we also found effects of *w*AlbB infection on adult heat tolerance^50^, but the effect was much smaller and only apparent in one genetic background. The exact reason for this difference is unclear, but could be due different backgrounds being tested or differences in mosquito age, given that *w*AlbB *Wolbachia* density increases with adult age as shown here and high densities are associated with stronger fitness costs^48^. Consistent with our previous work^41^, we also found costs of *Wolbachia* infection when eggs were heat-stressed. However, it is unclear if *Wolbachia* reduces thermal tolerance when there are also general fitness costs, especially given that the slope of the decrease in hatch was similar between *Wolbachia*-infected and uninfected lines. Our experiments showing substantial costs of *w*AlbB infection after storage under different temperature and RH conditions also support the idea that the costs of *Wolbachia* infection are not strongly temperature-dependent.

Dengue hemorrhagic fever has been reported in Saudi Arabia since 1994^80^ with cases fluctuating between 1500 up to 7000 cases spanning a decade from 2006 to 2016, resulting in an economic burden of US$110 million annually. The most common dengue virus serotype has been DENV2. Two more serotypes were reported (DENV1 and DENV3); however, there is no record of DENV4 among published serological studies^81^. The strong blockage of dengue virus (DENV2) by *Wolbachia* and particularly *w*MelM in the Jeddah genetic background provides confidence for mosquito releases aimed at population replacement to proceed in the Jeddah area. At present, dengue suppression in Jeddah is reliant on regular and widespread applications of pesticides, which is not sustainable in the long run due to both environmental costs and the evolution of resistance^57, 62^. It is still unclear how easily *Wolbachia* invasion will proceed within the Jeddah environment, and whether the replacement process can be facilitated through the selection of a particular *Wolbachia* strain and releases targeting a particular time of the year. Nevertheless, even partial replacement is likely to have beneficial outcomes in terms of dengue suppression^19, 34^, and there is clearly the added benefit of fitness costs incurred by the *Wolbachia* strains leading to a reduction in mosquito abundance which can benefit both dengue transmission^82^ and nuisance mosquito biting, potentially reducing the pesticide load in urban environments.

## 4. Materials and methods

### 4.1 Mosquito strains and colony maintenance

We performed experiments with laboratory populations of *Ae. aegypti* mosquitoes from two geographic origins (Cairns, Australia and Jeddah, Saudi Arabia) and three *Wolbachia* infection types (*w*AlbB, *w*MelM and uninfected). The *w*AlbB infection was originally transferred from *Ae. albopictus* via microinjection^83^ and is a *w*AlbB-I variant of the strain^84^. The *w*MelM infection is a clade I variant of *w*Mel that was transferred from temperate *D. melanogaster* via microinjection^64^. *Wolbachia* infections were introduced to *Ae. aegypti* with an Australian mitochondrial haplotype and nuclear background through microinjection as described previously^63, 64^. *w*AlbB*-*infected mosquitoes were then treated with tetracycline^63^ to generate uninfected mosquitoes with matching genetics. To generate populations with homogenized Australian or Saudi Arabian backgrounds, females of each *Wolbachia* infection type were crossed to natively uninfected populations collected from Cairns, Australia in February 2018 or Jeddah, Saudi Arabia in October 2019. Females were backcrossed to Saudi Arabian males for four consecutive generations, resulting in an estimated 93.75% similarity to the target background. The resulting populations were named *w*AlbB Au, *w*MelM Au, Uninfected Au (Australian background) and *w*AlbB S, *w*MelM S, Uninfected S (Saudi Arabian background). Due to an initial focus on *w*AlbB strain for field release, all experiments involved the *w*AlbB infection while *w*MelM-infected mosquitoes were only included in a subset of experiments.

*Aedes aegypti* mosquitoes were maintained at 26 ± 1°C with a 12-h photoperiod and all experiments were performed under these conditions unless otherwise stated. *Wolbachia* genome sequencing, *Wolbachia* proliferation and experiments testing the effects of temperature and humidity on egg hatch were performed at the University of Cambridge, United Kingdom. Virus challenge experiments were performed under biosafety level 3 (BSL-3) conditions in an insectary at the Australian Centre of Disease Preparedness. All other experiments were performed in a quarantine insectary at the University of Melbourne, Australia. For experiments in Australia, populations were maintained at a census size of 400 individuals in 13.8 L BugDorm-4F2222 (Megaview Science C Ltd., Taichung, Taiwan) cages. Larvae were reared in trays filled with 4 L of reverse-osmosis (RO) water and provided with fish food (Hikari tropical sinking wafers; Kyorin Food, Himeji, Japan) *ad libitum* throughout their development. Females (5-7 d old, starved for 1 d) were fed human blood sourced from the Red Cross Lifeblood service (agreement numbers 16-10VIC-02 and 20-04VIC-09) via Hemotek membrane feeders (Hemotek Ltd., Blackburn Lancashire, Great Britain). For experiments in the United Kingdom, larvae were reared in pots (AzlonTM Straight-sided Beakers; 500mL) or trays (Addis Seal Tight rectangular; 3000mL) at controlled densities 100 or 200 larvae respectively and fed liver powder *ad libitum.* Adults were kept in 27 L BugDorm-1 cages (Megaview Science C Ltd., Taichung, Taiwan) and fed with a solution of 10% fructose and 0.1% para-aminobenzoic acid (PABA). Females were fed donated human blood from Blood Transfusion Services at Addenbrooke’s Hospital, Cambridge, UK via Hemotek membrane feeders. Eggs were hatched in reverse osmosis water with a small pinch of desiccated liver and a few grains of desiccated yeast.

### 4.2 Wolbachia *genome sequencing*

We sequenced the *Wolbachia* genome of the *w*AlbB S population and compared this to published sequences of *w*AlbB Au^63^ to investigate potential genetic changes following the introgression of the *w*AlbB infection into Saudi Arabian *Ae. aegypti.* Two DNA sequencing libraries were prepared from two *w*AlbB S mosquitoes using the NEBNext^®^ Ultra™ II FS DNA Library Prep Kit for Illumina (New England Biolabs, E7805L) and NEBNext^®^ Multiplex Oligos for Illumina^®^ (New England Biolabs, E7335G) according to the manufacturers’ recommendations with variations as follows: all samples were fragmented for 10 minutes at 37°C. A post-ligation size selection step was performed on both libraries, 25ul of KAPA Hyperpure beads (Roche 8963835001) were used for the first step and 10ul of beads for the second step. 7 cycles of post-adaptor ligation PCR enrichment were used. The libraries were quantified using a Qubit HS DNA assay kit (Thermofisher Q32851) and then combined to give a final quantity of 950ng in a volume of 45ul. Samples were enriched for *Wolbachia* target DNA using a KAPA Hypercapture kit (Roche 09075810001) and KAPA Hypercapture Bead kit (Roche 09075780001) with custom designed KAPA Hyperexplore Max probes (Roche 09063633001) according to the manufacturer’s recommendations. Alternative options in the protocol were available, where this was the case, 20ul of COT human DNA (1mg/ml) was used to block the hybridization reaction and 10 cycles of post-capture PCR amplification was used. Captures were eluted from the final bead wash in 20ul of EB, quantified using Qubit HS DNA assay kit and analyzed using a Tapestation D1000 kit (Agilent 5067-5582). Sequencing was performed using Illumina MiSeq with 2 x 250bp reads (University of Cambridge, Department of Biochemistry sequencing service). Sequences were submitted to the NCBI Sequence Read Archive under BioProject ID PRJNA857586.

### 4.3 Processing of sequencing data

As preliminary analysis revealed that the two replicate *w*AlbB Au libraries were identical, they were combined before analysis. We also reanalysed raw sequence reads of *w*AlbB Au^63^. AdapterRemoval was used to trim adapters sequences from paired-end FASTQ reads, which also merged the overlapping reads using collapse option. The merged and unmerged sequence reads were aligned separately to the *Wolbachia pipientis w*AlbB chromosome (GenBank accession no. CP031221.1) using BWA-MEM with default options. BAM files belonging to the same sample were then merged using SAMtools. MarkDuplicates module in Picard was used to remove PCR and duplicates from the BAM files. Reads with a mapping quality of <30 were removed from the alignment, and coverage metrics were subsequently calculated using module DepthOfCoverage in GATK. Variant calling was performed for each individual using the HaplotypeCaller tool from GATK, and genotyping was then conducted using tool GenotypeGVCFs. The raw variant calling files were filtered by the set of hard filters to remove potential false variants, using the standard parameters recommended by GATK.

### 4.4 Cytoplasmic incompatibility

We performed reciprocal crosses to test the ability of *w*AlbB to induce cytoplasmic incompatibility in *Ae. aegypti* with a Saudi Arabian background. *w*AlbB S and Uninfected S males were crossed to *w*AlbB S or Uninfected S females (both 2 d old) and females were blood fed at 5 d old after being sugar-starved for 24 hr. Twenty females from each cross were isolated for oviposition in 70 mL specimen cups with mesh lids lined with sandpaper strips and filled with 15 mL of larval rearing water. Eggs were collected 4 d after blood feeding, partially dried, then hatched 3 d post-collection. Egg hatch proportions were determined 3 d after hatching by counting the number of hatched (egg cap detached) and unhatched (intact) eggs.

### 4.5 Wolbachia maternal transmission

We assessed the fidelity of *w*AlbB and *w*MelM maternal transmission in *Ae. aegypti* with a Saudi Arabian background. Females from the *w*AlbB S and *w*MelM S populations were crossed to Uninfected S males and fifteen females per population were blood fed and isolated for oviposition. We then screened ten offspring from ten females (110 individuals per population in total) for the presence of *Wolbachia* (see below). In a second experiment, we measured the maternal transmission of *w*AlbB S infection at a later date by testing 75 female offspring from 17 female parents.

### 4.6 *wAlbB* Wolbachia *proliferation in adult mosquitoes*

We investigated the proliferation of the *w*AlbB infection in adult female mosquitoes with a Saudi Arabian background. Larvae were reared to the pupal stage and separated by sex. Virgin *w*AlbB S females were fed on fructose and PABA, then collected at days 0, 7, 14, 21, 28 and 35 days post-eclosion. We then measured *w*AlbB densities in 40 females from each time point across two independent experimental replicates (see *Wolbachia quantification*).

### 4.7 DENV2 virus challenge

We tested the effect of *w*AlbB and *w*MelM strains on dengue virus serotype 2 (DENV2) infection in Saudi Arabian *Ae. aegypti* through an artificial membrane feeding assay^85^. DENV2 isolate ET300 (MT921572.1) was passaged in Vero cells and viral supernatant was used in experiments. The final virus stock titer was 10^6^ TCID50/ml in Vero cells. In the first experiment, we compared DENV2 infection frequencies and titers between uninfected, *w*AlbB-infected and *w*MelM-infected populations with a Saudi Arabian background (Uninfected S, *w*AlbB S and *w*MelM S). In a second experiment, we compared uninfected and *w*AlbB-infected populations with an Australian background (Uninfected Au and *w*AlbB Au). In a third experiment, we performed a repeat of the Uninfected S and *w*MelM S comparison.

Female mosquitoes (6-12 d old) were artificially fed a meal of human blood (agreement number 20-04VIC-09) for 30 mins using a collagen membrane and a Hemotek membrane feeding system. For each infection, human blood was centrifuged at 2500 rpm for three minutes on a tabletop centrifuge, plasma discarded, then DENV2 viral supernatant was mixed with an equal volume of red blood cells for a final DENV2 titer of 10^6^ TCID50/ml. Mosquitoes were then chilled and anesthetized with carbon dioxide. Engorged females were transferred to plexiglass cages and housed at 27°C, 65% relative humidity with a 12:12 hr light/dark cycle and provided with 10% sugar ad libitum and an oviposition substrate.

### 4.8 DENV2 quantification

Samples from blood-fed female mosquitoes were collected as previously described^86^. Quantitative PCR (qPCR) reactions were prepared in triplicate using TB Green^®^ Premix Ex Taq™ II (Tli RNase H Plus) (Takara, USA) with a modified protocol. Each reaction consisted of 2μL cDNA, 10μL TB Green Premix Ex Taq II, 0.4μL ROX, 0.4μL 10μM forward primer, 0.4μL 10μM reverse primer and 6.8μL nuclease free water in a total volume of 20μL. Quantitative PCR was performed on QuantStudio6 Flex PCR System (Applied Biosystems, USA) using DENV2 non-structural protein 1 primers (Table S2) with the following cycle parameters – Initial denaturation at 95°C for 30 seconds, 40 cycles at 95°C for 5 seconds, 60 °C for 30 seconds plus melt curve. *Ae. aegypti* Ribosomal protein S17 (RpS17) were used to normalize total RNA. After each run, threshold was set to 0.04 with an automatic baseline and data exported for detailed analysis.

To determine the absolute number of DENV2 genomic copies in each mosquito sample, a recombinant plasmids containing DENV2 ET300 fragment (2489bp – 2598bp) or *Aedes aegypti* RpS17 fragment (228bp-295bp) (Integrated DNA Technologies, Singapore) were used to generate a standard curve. Briefly, serial 10-fold dilutions of each plasmid were prepared in 10mM TE buffer with 50ng/μL yeast RNA (Sigma, USA) and each dilution tested via qPCR as previously described. Limits of detection (LOD) was 2 plasmid copies per reaction. Linear equations were derived from the standard curve, enabling normalized DENV2 genomic copy numbers to be calculated from C_T_ values obtained in qPCR of a DENV2 infected mosquito sample. For calculation, samples with a negative log value were interpreted as below LOD and were assigned a value of 0.

### 4.9 Larval development and body size

We measured larval development time, survival to pupa and adult wing length for each population. Eggs (< 1 week old) were hatched in plastic trays filled with RO water and a few grains of yeast. Larvae (< 8 hr old) were then counted into 750 mL plastic trays filled with 500 mL RO water and provided with fish food *ad libitum* throughout their development. We set up six replicate trays with 100 larvae per tray for each population. Pupae were counted, sexed and removed from trays twice per day. Adults emerged in cages and twenty adults of each sex were then measured for their wing length according to previous methods^51^.

### 4.10 Female fertility following egg storage

We measured the viability of quiescent eggs stored for up to 11 weeks and the fertility of females hatching from these eggs. Eggs were collected on sandpaper strips from colony cages of *w*AlbB Au, *w*AlbB S, Uninfected Au and Uninfected S populations. Sandpaper strips were partially dried for 2 d then placed in a sealed chamber with a saturated solution of potassium chloride to maintain relative humidity (RH) at ~80%. Eggs were then hatched after 1, 3, 5, 7, 9 and 11 weeks of storage by submerging sandpaper strips in trays of RO water with a few grains of yeast. Twelve replicate batches of eggs (range: 39-129 eggs per replicate) were hatched per time point per population. Egg hatch proportions were determined 3 d after hatching as described above. Larvae were controlled to a density of 50 larvae in 500 mL RO water and reared to adulthood. We then assessed fertility in females hatching from stored eggs by blood feeding and isolating 50 females per population at each time point. We scored the proportion of infertile females (that did not lay eggs after one week) and the number of eggs laid by fertile females and their hatch proportions. Females that died before egg laying were excluded from the analysis. Female fertility was not measured after 11 weeks of storage due to low egg viability in some populations.

### 4.11 Adult desiccation tolerance

We measured the survival of adult *Ae. aegypti* from the *w*AlbB Au, Uninfected Au, *w*AlbB S and Uninfected S populations under low (~15%) and high (~80%) RH as an estimate of desiccation tolerance. In each experiment, six (low RH) or eight (high RH) adults (5 d old, sugar-starved for 1 d) were aspirated into numbered glass vials with mesh lids, with eight replicate vials per sex per population. Vials were placed in sealed chambers in a random order around the perimeter so that mosquitoes could be observed without opening the chamber. We determined mortality by the tapping chamber and observing movement; mosquitoes that did not move after 10 s were considered dead. In the low RH experiment, mortality was scored every two hr, while in the remaining trials mortality was scored two to three times per day (four hr apart). In the low RH experiment, vials were placed in a sealed chamber containing trays with silica beads which maintained RH at ~15% within 3 hr of sealing the chamber. In the high RH experiment, containers filled with a saturated solution of potassium chloride were placed in the chamber to maintain RH at ~80%. Temperature and RH in each experiment was monitored with data loggers (Thermochron).

### 4.12 Adult water loss

We estimated water loss in adults from the *w*AlbB Au, Uninfected Au, *w*AlbB S and Uninfected S populations using methods adapted from previous studies^87, 88^. Adults (5 d old, sugar-starved for 1 d) were aspirated into numbered glass vials covered with a mesh lid, with eight adults per vial and eight replicate vials per sex per population. Vials were weighed on an analytical balance (Sartorius BP 210 D) then placed in a sealed chamber under conditions identical to the adult desiccation experiments at 15% RH. After 8 hr, vials were removed from the chamber and immediately weighed again. No mortality was observed during this period. Mosquitoes were killed by placing vials at −20°C for 48 hr then dried by placing vials in an oven at 70°C for 3 d. Vials were weighed before and after removing the dried mosquitoes. Fresh mass per mosquito was calculated by subtracting the empty vial mass from the mass of vials containing mosquitoes prior to desiccation, divided by eight. Dry mass per mosquito was subtracted was calculated by subtracting the empty vial mass from the mass of vials containing dried mosquitoes, divided by eight. Water mass was calculated by subtracting dry mass from fresh mass, while percent water content was calculated by dividing water mass by fresh mass. Percent water loss per hr was calculated by subtracting the mass of vials after 8 hr at 15% RH from the mass of vials containing mosquitoes prior to desiccation, then dividing by eight. Then, this value was divided by water mass and multiplied by 100. Two vials were excluded from the analysis due to escapees during the experiment.

### 4.13 Adult thermal tolerance

We measured the thermal tolerance of adult female *Ae. aegypti* from the *w*AlbB Au, Uninfected Au, *w*AlbB S and Uninfected S populations in a static knockdown assay based on a previous study ^50^. Female adults (5 d old, mated, starved for 1 d) were aspirated into numbered glass vials (50mm height x 18 mm diameter, 10mL volume), sealed with a plastic cap and fastened to a plastic rack with elastic in a randomized order. The rack was submerged in a water tank heated to a constant 42°C (Ratek Instruments, Boronia, Victoria, Australia). Mosquitoes in vials were observed and the time until knockdown (where mosquitoes were unable to stand up after vials were tapped with a metal rod) was recorded. Each trial involved 10 adult females per population. Trials were performed six times on separate days for a total of 60 replicates per population. Water temperatures were recorded using data loggers (Thermochron; 1-Wire, iButton.com, Dallas Semiconductors, Sunnyvale, CA, USA) placed in a glass vial that was held in the center of the plastic rack.

### 4.14 Egg viability and Wolbachia densities following exposure to cyclical heat stress

We measured egg hatch and *Wolbachia* titers at high temperatures by hatching eggs from the *w*AlbB Au, *w*MelM Au, Uninfected Au, *w*AlbB S, *w*MelM S and Uninfected S populations following exposure to a range of cyclical temperature regimes. Eggs from each population were collected on filter paper (Whatman 125 mm qualitative circles, GE Healthcare Australia Pty. Ltd., Parramatta, New South Wales, Australia), partially dried, then stored for 3 d at room temperature. Batches of eggs (range: 15-60) in 0.2 mL tubes were placed in blocks of TProfessional TRIO 48 thermocyclers (Biometra, Gottingen, Germany). Eggs were exposed to a constant 26°C or daily cycles of 27-37°C, 28-38°C, 29-39°C, 30-40°C, 31-41°C or 32-42°C according to methods described previously^41, 89^. Between 12 and 20 replicate tubes per population were exposed to each temperature cycle; low numbers of eggs laid by some populations resulted in variable numbers of replicates. Eggs were hatched after one week of exposure by brushing eggs from each replicate tube into wells of Costar 12-well cell culture plates (Corning, NY) filled with 4 mL of RO water and a few grains of yeast. Egg hatch proportions were determined one week after hatching according to methods described in the section above. Larvae hatching from the *Wolbachia*-infected populations at each temperature were pooled across replicates and reared at a density of 50 larvae in trays of 500 mL water. Adults were stored in absolute ethanol within 24 hr of emergence to measure *w*AlbB and *w*MelM and densities (see *Wolbachia quantification).* Nine adults per sex, per population, per temperature cycle were screened for *Wolbachia* density. Adults from the 31-41°C or 32-42°C treatments were not screened due to near-zero hatch proportions under these conditions.

### 4.15 Egg viability under different temperature and humidity conditions

We tested the effects of *w*AlbB infection on the hatch proportion of stored eggs under different temperature and RH conditions in two experiments. Blood-fed females from the Uninfected S and *w*AlbB S populations were transferred to cups in groups of five and eggs were collected every two days over a six day period. Each collection of eggs was divided evenly between the experimental treatments by cutting the paper. We hatched eggs at 48-72 hours after collection (‘control’; 80% RH and 26°C) or 21 days after the beginning of the experiment for each treatment under different temperature and humidity conditions. In the first experiment, eggs were stored at cycling temperatures of 20-30, 23-33, 26-36 or 29-39°C and 80% RH, with 12 hr at each temperature. In the second experiment, eggs were stored at 20-30°C or 29-39°C and 80% or 30% RH. To control RH, the eggs were placed in a sealed plastic box that contained an open beaker of a saturated hygroscopic salt solution. Saturated magnesium chloride hexahydrate (MgCl2·6H2O) solution maintained RH at ~30% while saturated potassium chloride (KCl) solution maintained RH at ~80%. We monitored the temperature and humidity in all the eggs storage boxes using a data logger to ensure these treatments were successful. Egg hatch proportions were determined for 15-25 replicate batches of eggs per treatment by counting the number of eggs and larvae 4-5 days after hatching.

### 4.16 Wolbachia detection and quantification

*Aedes aegypti* from colonies and experiments were screened to confirm their *Wolbachia* infection status and estimate relative *Wolbachia* density. We confirmed the presence or absence of *Wolbachia* in mothers and offspring from the maternal transmission experiment and screened colonies used in experiments routinely (approximately every 3 months). We also quantified *Wolbachia* densities in adults of different ages and in adults following exposure of eggs to cyclical heat stress.

For experiments in Australia, genomic DNA was extracted using 250 mL of 5% Chelex 100 resin (Bio-Rad Laboratories, Hercules, CA) and 3 mL of proteinase K (20 mg/mL) (Roche Diagnostics Australia Pty. Ltd., Castle Hill, New South Wales, Australia). Tubes were incubated for 60 min at 65°C and then for 10 min at 90°C. We ran real-time PCR assays ^90, 91^ using a Roche LightCycler 480 using primer sets to amplify an *Ae. aegypti*-specific marker (aRpS6_F [5’-ATCAAGAAGCGCCGTGTCG-3’] and aRpS6_R [5’-CAGGTGCAGGATCTTCATGTATTCG-3’]) and a *Wolbachia*-specific marker (*w*M*w*A_F [5’-GAAGTTGAAGCACAGTGTACCTT-3’] and *w*M*w*A_R [5’-GCTTGATATTCCTGTAGATTCATC-3’]) that can distinguish between *w*AlbB and *w*MelM infections based on differences in melting temperature (80.4 ± 0.02°C and 82.6 ± 0.03°C and respectively). Relative *Wolbachia* densities were determined by subtracting the *w*M*w*A crossing point (Cp) value from the aRpS6 Cp value. Differences in Cp values were averaged across 2 or more consistent replicate runs and then transformed by 2n.

For experiments in the United Kingdom, mosquitoes were homogenized in 1.7 mL tubes with 50 μL of water and approximately ten 1.0 mm diameter zirconia beads (Thistle Scientific 11079110zx) by shaking at 30 Hz for 2 minutes in a Tissue Lyzer II homogenizer (Qiagen). DNA was extracted using a PureLink Genomic DNA Mini kit (Thermofisher Scientific, K1820-02) by adding 180 μL of PureLink^®^ Genomic Digestion Buffer and 20 μL of Proteinase K solution, followed by mixing and incubation at 55°C for 2 hours. Samples were centrifuged at 13,000 g for 5 minutes to pellet debris and the supernatant was removed to a fresh tube. 400 μL of a 1:1 mixture of PureLink^®^ Genomic Lysis/Binding Buffer: Ethanol was added, and samples were mixed well by vortexing. Genomic DNA was purified using the standard spin column protocol according to the manufacturers recommendations and eluted from the column with 50ul PureLink^®^ Genomic Elution Buffer.

Quantitative PCR was carried out using Hi-Rox SensiFAST Sybr reagent (Meridian BIO-92020). For each qPCR plate a master mix was made containing 5 μL of SensiFAST master mix, 2 μL of nuclease free water and 1 μL of 2.5uM of each primer per reaction. The primers to amplify *w*AlbB were qwAlbB_Fw (5’-CCTTACCTCCTGCACAACAA) and qwAlbB_Rv (5’-GGATTGTCCAGTGGCCTTA)^42^. The primers to amplify mosquito DNA were mRpS6_Fw (5’-AGTTGAACGTATCGTTTCCCGCTAC) and mRpS6_Rv (5’-GAAGTGACGCAGCTTGTGGTCGTCC)^42^. After adding 8 μL of the master mix to wells of a 384-well MicroAmp Optical Reaction Plate (Applied Biosystems 4309849), 2 μL of genomic DNA sample which had been diluted 2:1 with nuclease free water was added. Plates were sealed using MicroAmp Optical Adhesive film (Applied Biosystems 4311971), vortexed to mix, and centrifuged briefly. The plates were run on a 384-well QuantStudio™ 5 Real-Time PCR System (Applied Biosystems A28575) using the pre-set comparative-CT-Sybr with melt curve program with the reaction volume adjusted to 10 μL. Data were analyzed using QuantStudio™ Design and Analysis Software v1.5.2. The CT threshold value was set to 0.1 for each plate.

### 4.17 Statistical analysis

All statistical analyses were performed in SPSS Statistics 26. Proportional data such as egg hatch were logit transformed before analysis ^92^, treating each replicate batch of eggs (or emerged adults) as an independent data point rather than each egg given that different eggs in a batch may have come from the same female and that each batch may have experienced slightly different exposure conditions. *Wolbachia* density data were analyzed prior to 2^n^ transformation. We ran general linear models for each trait, with genetic background and *Wolbachia* infection included as factors. Where multiple traits were extracted from the same data set, P values were corrected by the Bonferroni method based on the number of traits scored. When both sexes were tested, data for each sex were analyzed separately. For the adult heat tolerance experiment, we performed an analysis with a trial by population interaction. Because there were apparent differences in the relative performance of populations across trials, we reanalyzed data separately for each trial. Adult survival at low and high relative humidity was analyzed with Cox regression, with genetic background and *Wolbachia* infection included as factors. In experiments with multiple time points or temperature treatments, we analyzed data separately for each time point/treatment. For analyses of *Wolbachia* density, we included temperature, *Wolbachia* strain, background and their interactions as factors. For comparisons of female infertility, we computed 95% binomial confidence intervals on the proportion of females that did not lay eggs using the Wilson method ^93^. We compared DENV virus copy numbers between populations using non-parametric Mann-Whitney U tests, with negative samples excluded from the analysis. We also compared DENV2 infection frequencies (i.e. the number of samples testing positive for DENV2 out of the total number tested) between populations with two-tailed Fisher’s tests.

## Acknowledgements

We thank the municipality of Jeddah, Saudi Arabia for supporting this research study and facilitating fieldwork and sampling operations which allowed us to have access and build wild mosquito colonies. We thank Nancy Endersby-Harshman for facilitating the import of mosquito populations into the quarantine insectary. We thank Meng-Jia Lau for advice on primers and Véronique Paris for assistance with colony maintenance. We thank Kim R Blasdell for providing lab assistance at ACDP.

## Author contributions

PAR contributed to study design, experimental work, analysis, interpretation, and drafting the paper. SE contributed to experimental work, interpretation, funding, local resources, and drafting of the paper. SC, JKA and XG contributed to the experimental work on *Wolbachia* host comparisons. MJK, MD, SJ and PNP contributed to the design, analysis, experimental work and drafting of the vector competence component. SSL, YZ and FMJ contributed to *Wolbachia* host comparisons and *Wolbachia* sequencing as well as the drafting of the sequencing component. AAH contributed to interpretation, overall design, funding, and drafting of the paper. MBA-F, AMA, MSN and EAT contributed to project design, funding, supervising, local resources, as well as substantive revisions to the paper. All authors edited and approved the final version of the manuscript.

## Data availability

Raw sequence reads for the *Wolbachia* genomes in this study have been deposited in the NCBI Sequence Read Archive under BioProject ID PRJNA857586. All other data are contained within the manuscript and its supporting information files.

## Funding

This work was funded under the KACST-CSIRO co-investment collaborative research agreement [ETSC&KACST-CSIRO-2018-12-30-21] on “Management strategies of vector-borne disease in Saudi Arabia: feasibility of the *Wolbachia-based* approach as an alternative to chemical pesticides.” SE was sponsored by the CSIRO Julius career award (WBS: R-91040-11); AAH was partly supported by the National Health and Medical Research Council (1132412, 1118640, www.nhmrc.gov.au).

## Supplementary information

**Figure S1.**
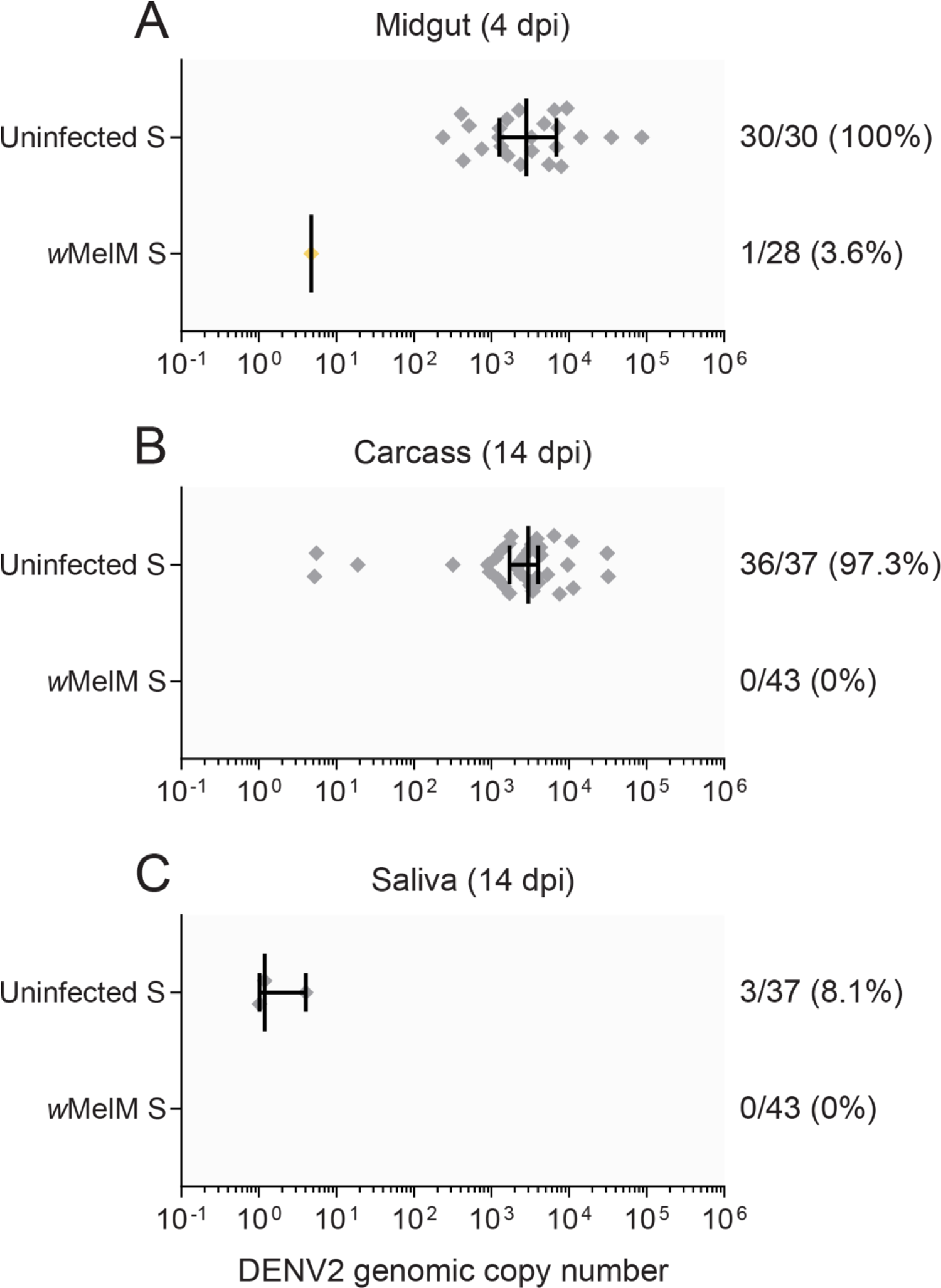
Comparison of DENV2 infection frequencies and copy numbers in uninfected and *w*MelM-infected *Ae. aegypti* with a Saudi Arabian background. We measured (A) midgut infection at 4 dpi, (B) dissemination to the carcass at 14 dpi and (C) saliva infection at 14 dpi as a proxy for transmission. Each symbol shows data from a single DENV2-positive sample (negative samples were excluded), while vertical lines and error bars represent medians and 95% confidence intervals. Numbers to the right of indicate the number of individuals positive for DENV2 out of the total number tested, followed by the percent positive.

**Figure S2.**
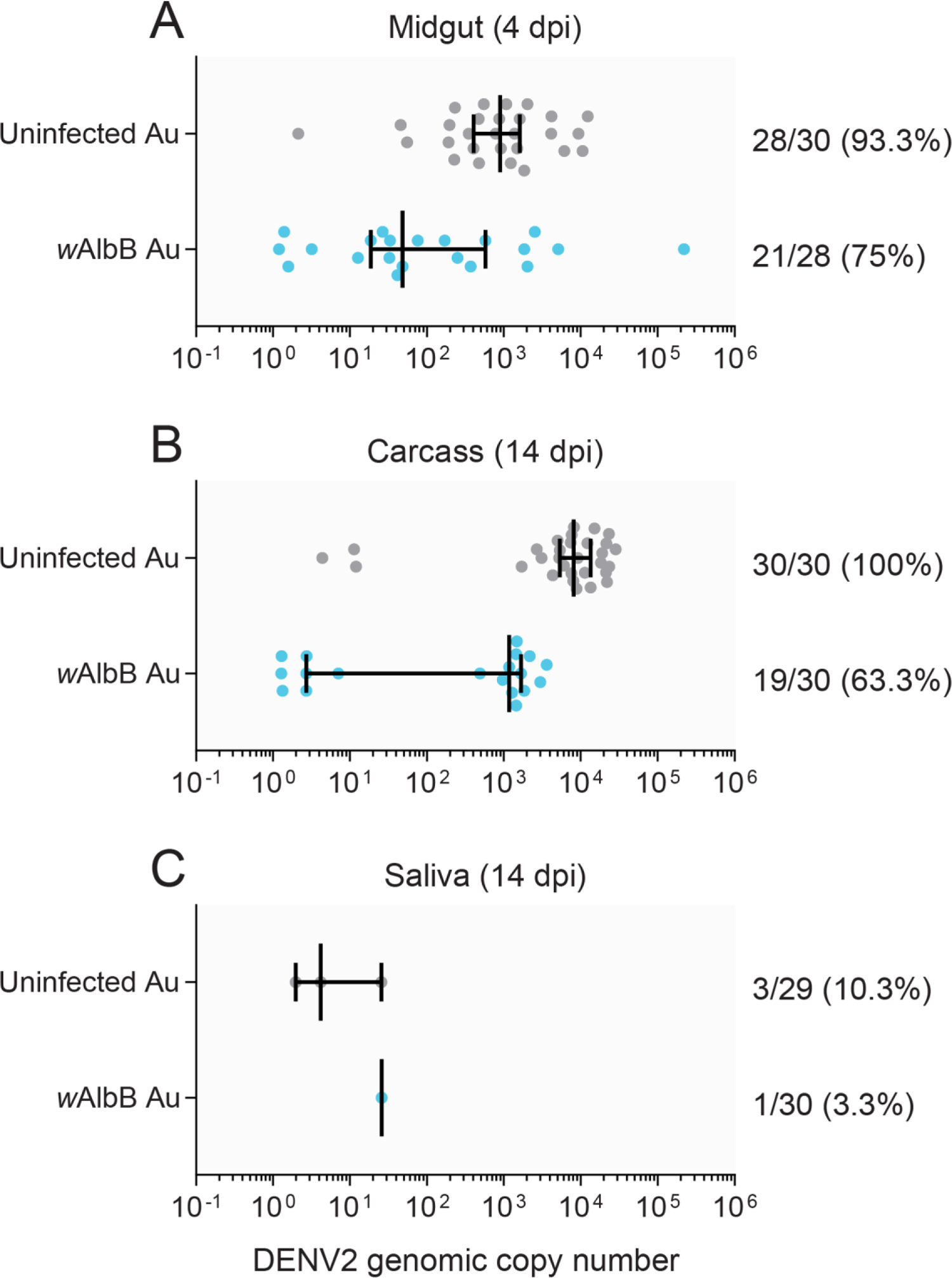
Comparison of DENV2 infection frequencies and copy numbers in uninfected and *w*AlbB-infected *Ae. aegypti* with an Australian background. We measured (A) midgut infection at 4 dpi, (B) dissemination to the carcass at 14 dpi and (C) saliva infection at 14 dpi as a proxy for transmission. Each symbol shows data from a single DENV2-positive sample (negative samples were excluded), while vertical lines and error bars represent medians and 95% confidence intervals. Numbers to the right of indicate the number of individuals positive for DENV2 out of the total number tested, followed by the percent positive.

**Figure S3.**
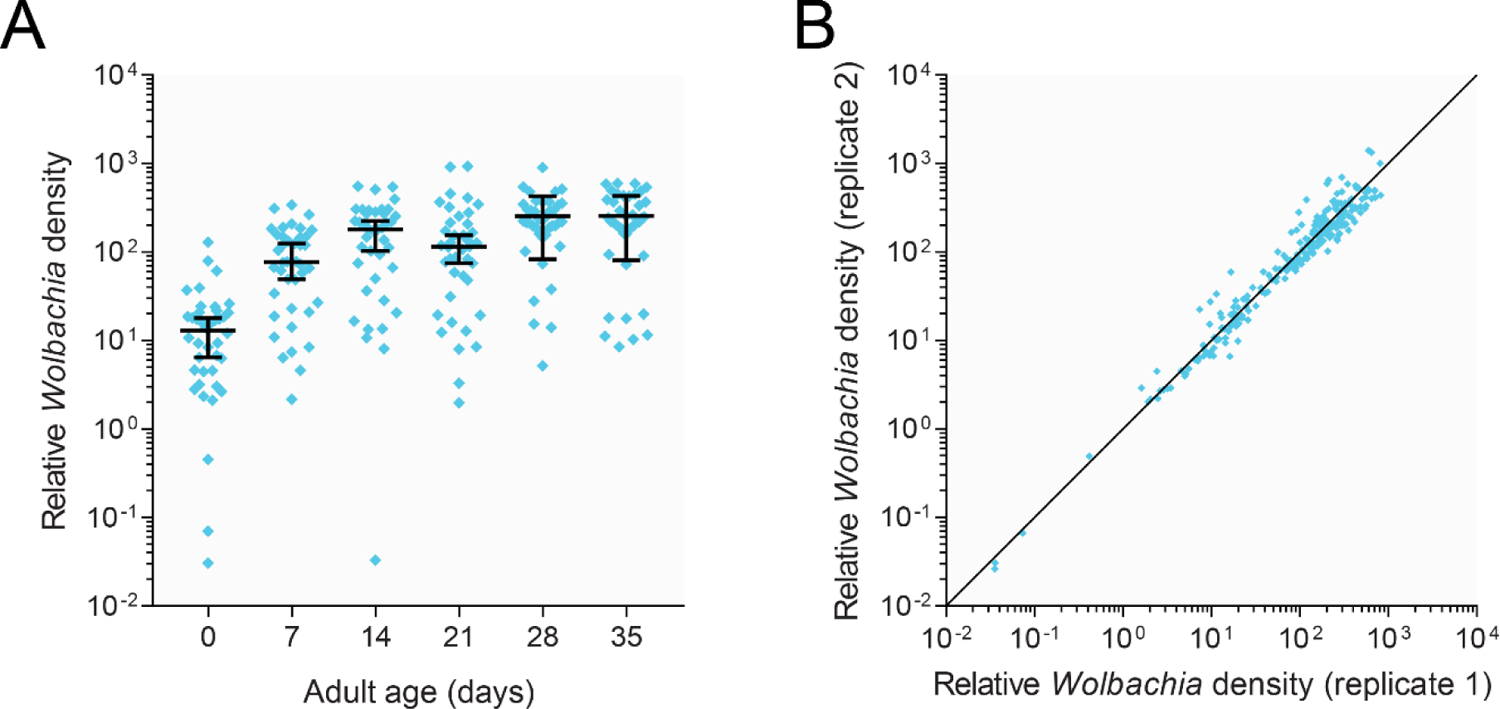
Changes in *Wolbachia* density with age in adult *w*AlbB S *Ae. aegypti* females. (A) *Wolbachia* density was measured every 7 days by quantitative PCR. Each point shows the relative amount of *Wolbachia* genomic DNA relative to mosquito genomic DNA in a single mosquito across two independent technical replicates. Horizontal lines and error bars represent medians and 95% confidence intervals. (B) Correlation in *Wolbachia* density between two replicate measurements on the same sample.

**Figure S4.**
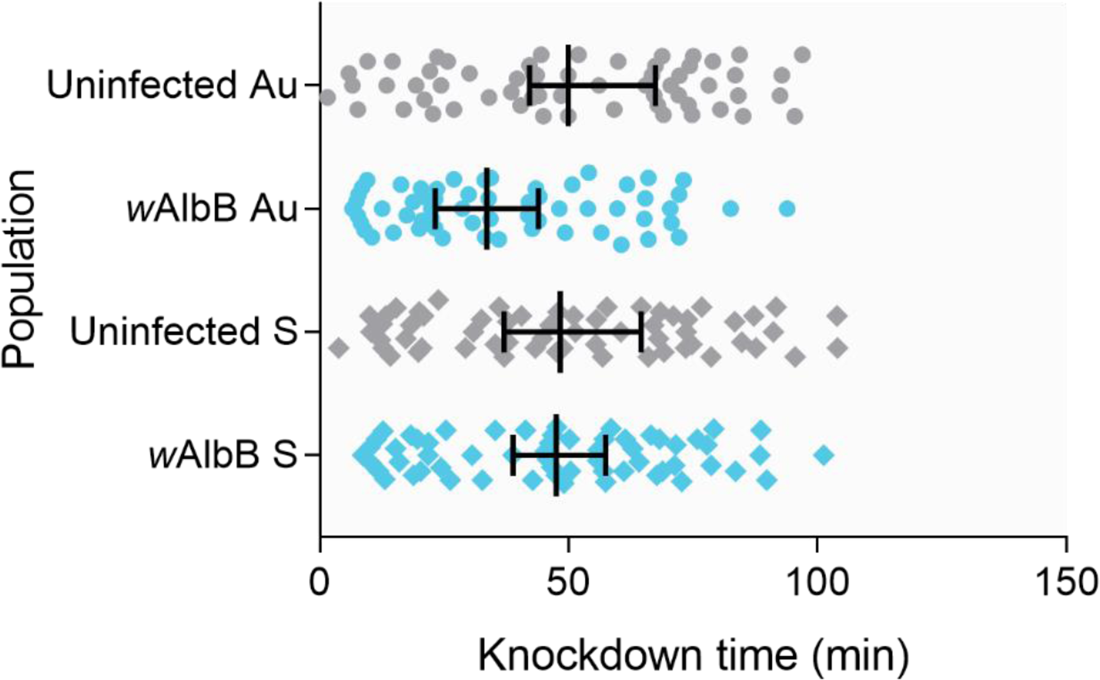
Knockdown time of *w*AlbB-infected and uninfected *Aedes aegypti* females from Australian and Saudi Arabian backgrounds exposed to a 42°C heat shock. Data are pooled across six replicate runs, with runs presented separately in the main text. Each symbol represents data from a single female, while vertical lines and error bars represent medians and 95% confidence intervals.

**Figure S5.**
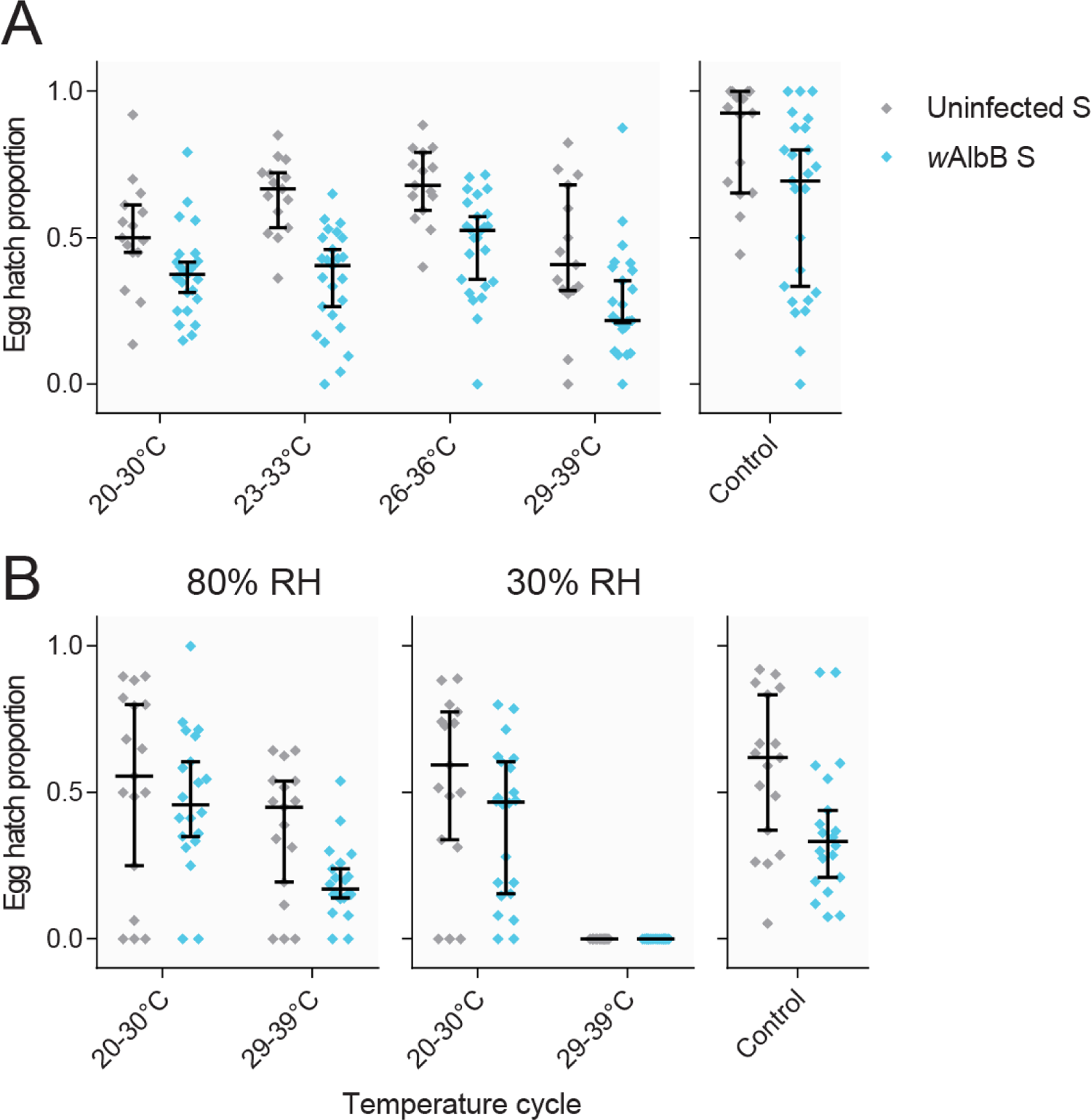
Effects of temperature, humidity and *w*AlbB infection on egg hatch following storage in Saudi Arabian *Ae. aegypti.* (A) Egg hatch proportions of *w*AlbB S and Uninfected S eggs stored for 21 days at four different temperature cycles (12h: 12h) at 80% relative humidity (RH). *w*AlbB infection reduced egg hatch across all temperature cycles (GLM: *Wolbachia* main effect: F_1,152_ = 32.887, P < 0.001) with no significant interaction between *w*AlbB infection and temperature cycle (F_3,152_ = 0.197, P = 0.313). (B) Egg hatch proportions of *w*AlbB S and Uninfected S eggs stored for 21 days at 80% or 30% RH and two different temperature cycles. In both experiments, control eggs were stored for 4-5 days at 26°C, 80% RH. All panels show combined data from two independent replicates of each experiment. Each symbol represents a replicate egg hatch measurement, while horizontal lines and error bars represent medians and 95% confidence intervals.

**Table S1.** Nucleotide differences between the *w*AlbB genomes sequenced in this study and the *w*AlbB reference genome.

| Position on genome | Reference allele | Alternate allele |
| --- | --- | --- |
| 404124 | A | G |
| 507205 | T | C |
| 542038 | G | A |
| 799342 | G | GA |
| 1373220 | C | T |

**Table S2.**
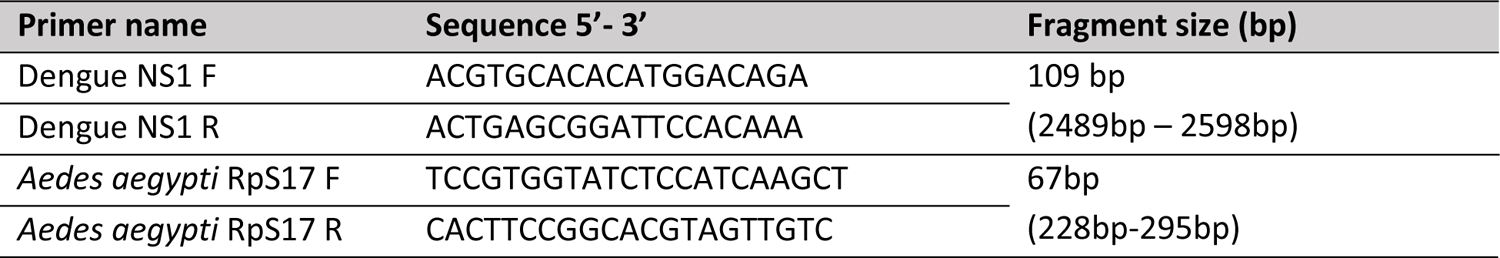
Primers used in qPCR for detection of DENV2 and *Aedes aegypti* RpS17 nucleic acid.

## Notes

### Competing Interest Statement

The authors have declared no competing interest.

### Summary of Updates

The manuscript has been updated to correct a spelling error in the author list

